# Prioritized docking of synaptic vesicles provided by a rapid recycling pathway

**DOI:** 10.1101/2022.06.08.495263

**Authors:** Van Tran, Alain Marty

## Abstract

It is known that endocytosis of synaptic vesicles, and docking of these vesicles to their release sites, are regulated in a similar manner, but it has remained unclear whether the two processes are linked together mechanistically. To address this issue, we studied vesicular release during repeated trains of presynaptic action potentials. Synaptic responses decreased when the inter-train interval was shortened, indicating a gradual exhaustion of the recycling pool of vesicles, with a resting capacity of about 180 vesicles per active zone. This effect was partially counteracted by activation of a rapid recycling pathway that depended on the activation of myosin light chain kinase and that had a capacity of 200 vesicles per active zone. Vesicles arising from the rapid recycling route were preferentially docked in comparison with vesicles coming from an upstream pool, demonstrating a differential sorting of vesicles inside the readily releasable pool depending on their origin.

## Introduction

Central synapses are challenged with potentially conflicting demands. On the one hand, they must transmit quickly and precisely information coming from trains of high-frequency presynaptic action potentials (APs). On the other hand, they must remain responsive even after several bouts of intense activity. Thus, synaptic vesicular release must combine high time precision, high reliability, and high resilience. A way that synapses use to manage these demands is to organize SVs into multiple functional pools (Rizzoli and Betz, 2005). The readily releasable pool (RRP) is responsible for the immediate release of neurotransmitter in response to AP stimulation. The RRP is replenished either by recycling fused SVs or by recruiting new SVs from an upstream pool. Collectively, all SVs that take part in synaptic transmission constitute the recycling pool. This pool typically contains 5-20% of all SVs within a synapse (Delgado et al., 2000; Harata et al., 2001; Lange et al., 2003; Richards et al., 2000, 2003; but see Ikeda and Bekkers, 2009; Xue et al., 2013). The remaining SVs, which form the reserve pool, only undergo exocytosis during intense and prolonged activity. These SVs have also been shown to act as buffers that bind to soluble synaptic proteins and prevent them from diffusing into the axon (Denker et al., 2011).

Vesicle recycling starts with the retrieval of fused vesicle membrane, followed by the reconstitution and refilling of SVs, and finally the recruitment of newly formed SVs to the RRP. At central mammalian synapses, endocytosis of SV membrane occurs in several kinetically different pathways, ranging from 50-100 ms (“ultrafast”; Watanabe et al., 2013) to seconds and tens of seconds. The contribution of these pathways depends on the stimulation intensity (Chanaday and Kavalali, 2018; Delvendahl et al., 2016; Midorikawa et al., 2014; Renden and Gersdorff, 2007; Soykan et al., 2017; Tanifuji et al., 2013; Wu et al., 2009; Yamashita et al., 2010). Once inside the synapse, recently endocytosed vesicles fuse to form endosome-like structures (Rizzoli et al., 2006), from which SVs are then regenerated in a clathrin-dependent manner (Watanabe et al., 2014). In an electron microscopy study at hippocampal synapses, newly formed SVs appeared ~3 s after ultrafast endocytosis of vesicle membrane (Watanabe et al., 2014). Importantly, some of these SVs were docked at the active zone membrane within 10 s after stimulation. The reacidification of SVs, which is required for the transport of neurotransmitter molecules into the vesicle lumen, has a time constant of ~5 s (Atluri and Ryan, 2006; Granseth et al., 2006). A similar time frame was observed for the refilling of SVs with glutamate at physiological temperature (time constant ~7 s; Hori and Takahashi, 2012). Altogether, these studies indicate a potential for glutamatergic synapses to reuse SVs every ~10 s. Faster recycling of SVs may be achieved by the kiss-and-run exo-endocytosis pathway where SVs recycle without passing through an endosome-like compartment (Fesce et al., 1994; Klingauf et al., 1998; Klyachko and Jackson, 2002; Pyle et al., 2000; Stevens and Williams, 2000).

Preferential recycling and reuse of recently fused SVs has been reported at hippocampal synapses (Ertunc et al., 2007; Pyle et al., 2000; Sara et al., 2002) and neuromuscular junctions (Maeno-Hikichi et al., 2011; Richards et al., 2003). At these synapses, recovery from synaptic depression depends on the recycling of RRP vesicles, rather than on the recruitment of vesicles from the reserve pool. However, the extent to which vesicle recycling contributes to RRP replenishment and prevents synaptic depression varies with the type of synapse (Gersdorff and Matthews, 1997) and the stimulation protocol (Ertunc et al., 2007; Fernández-Alfonso and Ryan, 2004). Recent studies further showed that the RRP is heterogenous, comprising several subpools that are differentially recruited and that have different roles during repetitive synaptic transmission (Miki et al., 2016; Neher and Brose, 2018; Neher and Taschenberger, 2021; Sakaba, 2006). It remains unclear through which subpools the recycled SVs enter the RRP.

Much knowledge has recently been obtained about vesicular docking, one of the last steps before exocytosis. Docking has been revealed as a fast (a few milliseconds only), reversible process that is enhanced by calcium and involves activation of the cytoskeleton (Chang et al., 2018; Kusick et al., 2020; Miki et al., 2016, 2018). Docking increasingly appears as a key control point of synaptic strength (Neher and Brose, 2018; Silva et al., 2021), but it remains unclear whether docking is regulated in conjunction with SV recycling.

In this study, we investigated whether glutamatergic synapses formed between cerebellar granule cells (GCs) and molecular layer interneurons (MLIs) preferentially recycle and reuse recently fused SVs. Similar to the majority of synapses in the brain, individual GC-MLI synapses typically contain one active zone (Masugi-Tokita et al., 2007; Xu-Friedman et al., 2001) and a relatively small number of SVs (Eguchi et al., 2020). However, compared to others, vesicular release statistics and SV pool dynamics have been better characterized at these synapses due to recent methodological advances (Malagon et al., 2016, 2020; Miki et al., 2016, 2018; Tran et al., 2022). Here, we found evidence for a rapid route of SV recycling during repeated train stimulation with short inter-train intervals. This route, which requires activation of the myosin light chain kinase, uses recently endocytosed vesicles to delay the reduction in synaptic output. It has a limited capacity and contributes minimally during steady-state release with short inter-train intervals. SVs that are recycled through this pathway are preferentially docked compared to those coming from the reserve pool. Prioritized docking thus reveals a functional link between a specific recycling pathway and SV sorting within the RRP. During and after extended synaptic stimulation, prioritized docking enhances the synaptic response to the first stimulation in a train, thus protecting responses to scarce stimuli from depression.

## Results

### Variation of synaptic response to short AP trains as a function of inter-train interval

In the present work, we studied the recycling of SVs during repetitive stimulation at cerebellar GC to MLI synapses. As in previous studies (Miki et al., 2017; Tran et al., 2022), we stimulated locally a single presynaptic GC and measured release at a single GC-MLI contact (**Fig. 1A**, left). Such contacts typically contain a single active zone (AZ) and are called for this reason “simple synapses”. As illustrated in **Fig. 1A** (right), simple GC-MLI synapses comprise several docking/release sites acting in parallel. These synapses have a large quantal size, so that the number of released SVs can be accurately determined by deconvolution of EPSC traces, using the mean quantal EPSC as template (Malagon et al., 2016). The number of released SVs following AP stimulation obeys a binomial distribution, where the binomial parameter N represents the number of docking/release sites (mean value of N per AZ: 4.0; Malagon et al., 2016). Analysis of cumulative SV number statistics indicates that docking is a two-step process (Miki et al., 2016). Each incoming SV transits through a replacement site before proceeding to an associated docking site (**Fig. 1A**, right).

**Figure 1:**
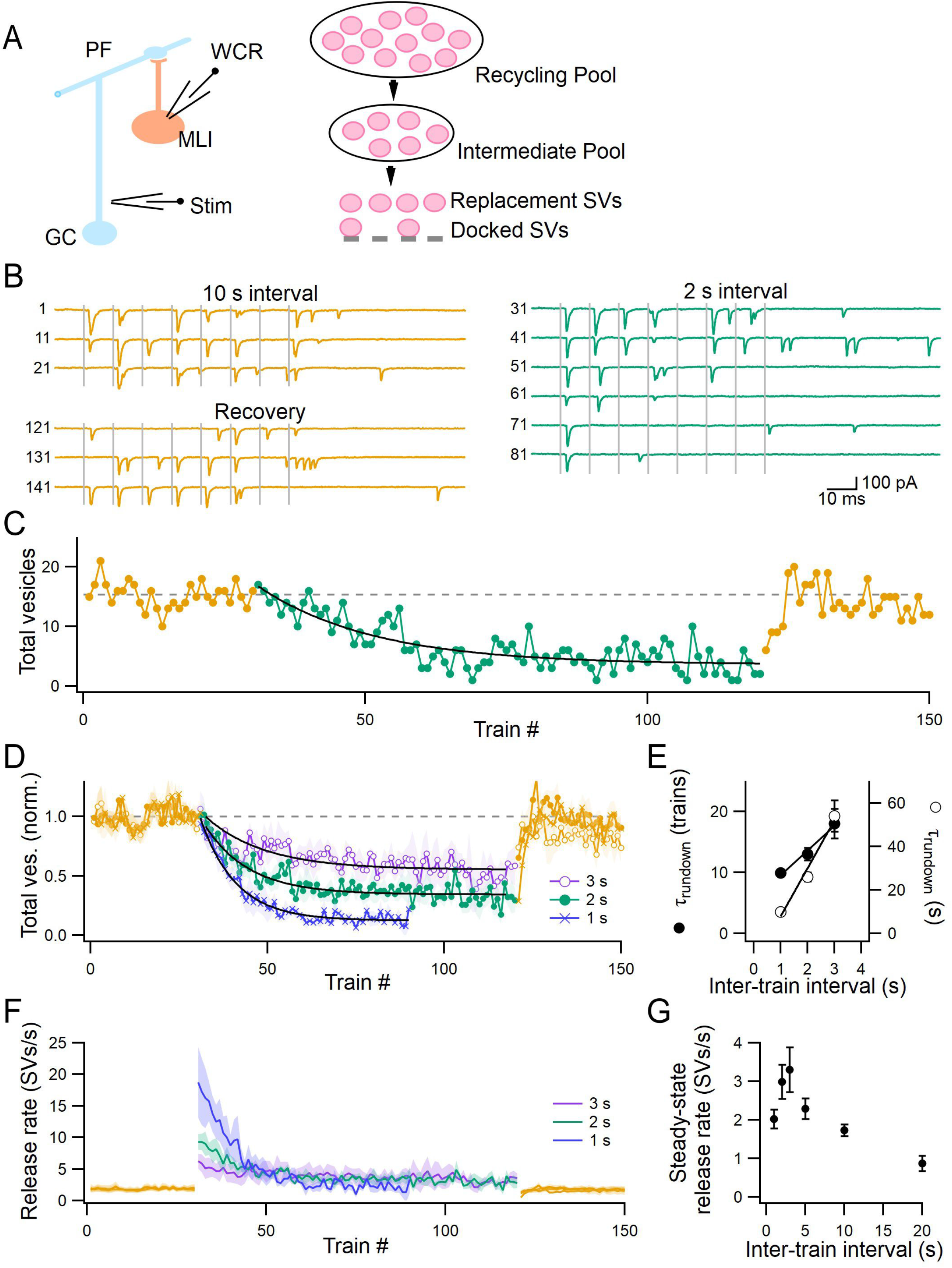
Evolution of summed SV numbers released by presynaptic AP trains during changes in the inter-train interval. **A, left**: Methodological approach. A whole-cell recording (WCR) was obtained in an MLI while a single presynaptic granule cell was being stimulated. **A, right**: Schematic of vesicular pools at a GC-MLI synapse. Each synapse contains a single AZ where several docking/release sites can release SVs in parallel. Each docking site is associated with a replacement site. Docking and replacement sites may or may not be occupied by a docked SV and a replacement SV, respectively. Replacement sites are replenished by a small-capacity intermediate pool, which is in turns replenished by the recycling pool. **B**: An example recording. Postsynaptic responses to 8-AP trains (100 Hz) were stable with 10 s inter-train intervals during the baseline period (upper left; train numbers as indicated). When the inter-train interval was reduced to 2 s, responses gradually decreased (right panel). Full recovery was obtained after the return of the inter-train interval to the initial value (lower left). **C**: Plot of total SV release as a function of train number for the same experiment. An exponential fit to the response during 2 s intervals yielded a time constant of 20 trains. **D**: Group data showing reversible reductions in cumulative SV release for 1-3 s inter-train intervals. Traces were fitted with exponential functions (black), the time constant of which are shown in E. **E**: The time constant of the reduction in total SV release per train, plotted against the inter-train interval. **F**: Group data showing the evolution of output frequency during changes in the inter-train interval. **G**: The steady-state output frequency plotted against the inter-train interval. n = 4-5 for each time interval. +/-SEM margins are shown as shaded areas in D and F, and as error bars in E and G.

To study the evolution of vesicular pools during repetitive synaptic activity, we applied trains of 8 presynaptic APs at 100 Hz under high release probability conditions (3 mM external calcium concentration, and 1 mM tetraethylammonium chloride or TEA). By changing the inter-train time interval in twin train experiments, we have recently shown that a single 8-AP train significantly reduces the size of an SV pool upstream of the RRP (called the intermediate pool or IP; **Fig. 1A**, right; Tran et al., 2022). Furthermore, we have shown that IP recovery occurs on a timescale of seconds, and is calcium-dependent (Tran et al., 2022). To explore further the nature and the size of SV pools upstream of the IP, we applied 8-AP trains repetitively, while changing the inter-train time interval from a baseline value of 10 s to various test values. In the example experiment of **Fig. 1B**, the synaptic response was stable with an inter-train interval of 10 s (top left; yellow traces). When switching to a 2 s interval, responses gradually became weaker as a function of train number (right; green traces), and they recovered upon returning to the 10 s interval (bottom left, yellow traces). In this example, the total number of SVs released per train decreased with a time constant of 20 trains (or 40 s) when the inter-train interval was reduced from 10 to 2 s, and it recovered to the baseline value within a few trains (corresponding to a few tens of seconds) when the interval was returned to 10 s **(Fig. 1C)**.

Group results show that, during prolonged train stimulation with short inter-train intervals (a total of 60 trains with 1 s interval or 90 trains with 2 or 3 s interval), the total number of SVs released per train gradually decreased before reaching a steady state (**Fig. 1D**). These reductions could therefore be approximated by single exponential functions, with time constants that increased as a function of the inter-train interval (from 10 trains or 10 s for 1 s interval to 18 trains or 54 s for 3 s interval: **Fig. 1E**). After the inter-train interval was returned to 10 s, the kinetics of recovery was rather poorly resolved due to the small number of test trains, but it was clear that responses returned to the baseline level within a few tens of seconds (**Fig. 1D**). Overall, changes in the total number of SVs released per train occurred over a time period of a few tens of seconds when the inter-train interval was altered.

The total number of released SVs was not changed when the inter-train interval was increased from 10 to 20 s (18 ± 4 SVs/train vs. 17 ± 4 SVs/train, respectively; p-value for paired t-test ppt = 0.5; n = 4; **Suppl. Fig. 1A**). This observation, together with the finding that the response with 10 s intervals was stable over time, indicates that the standard 10 s interval does not have a significant inhibitory effect on the response to 8-AP trains. On the other hand, the total release per train was significantly reduced during steady-state levels with short inter-train intervals (ppt < 0.05; n = 4-5 for each interval; **Suppl. Fig. 1A**). When release was separated into synchronous and asynchronous components, the relative reductions during steady-state levels were similar between these two release modes (**Suppl. Fig. 1B**).

With the standard 10 s inter-train interval, the SV output per train was ~17 SVs (including both synchronous and asynchronous release, up to 200 ms after stimulation onset), corresponding to an output frequency of ~1.7 SVs/s/AZ (the total number of SVs released per train divided by the inter-train interval). When the inter-train interval was decreased, the output frequency first increased to a level that was proportional to the increase in stimulation frequency (**Fig. 1F**). However, it soon declined to a steady-state level, with the 3 s interval yielding the largest steady-state output frequency (3.3 SVs/s/AZ or ~1.9-fold the baseline value; **Fig. 1G**). For intervals < 3 s, the steady-state output frequency was reduced to near baseline values (**Fig. 1G**). These results indicate different limits to the synaptic output frequency depending on the inter-train interval. Therefore, it is likely that multiple mechanisms underlie the steady-state SV release at short intervals.

In conclusion, the total number of SVs released per train during repetitive 8-AP train stimulation displays a gradual and reversible decrease for inter-train intervals of <10 s. Since the number of SVs in docking sites, replacement sites, and the IP, all replenish within 1-2 s after a single 8-AP train (Tran et al., 2022), the slow decrease observed upon train repetition likely reflects, at least in part, a gradual depletion of an SV pool located upstream of the IP.

### Differential depletion levels of docked, replacement, and intermediate SV pools with short inter-train intervals

To further investigate the mechanisms underlying the decrease in synaptic output at short inter-train intervals, we analyzed changes in the pattern of responses to individual APs. Examination of individual experiments suggested that the inhibition of synaptic output was associated with increased synaptic depression (green traces in **Fig. 1B**). This observation is paradoxical since in many cases of synaptic plasticity associated with presynaptic modifications, decreases in responsiveness are less prominent for late stimuli. In particular, a standard finding is that the paired-pulse ratio (PPR) increases while the response to the first stimulus (s_1_) decreases, as a decreased release probability both reduces s_1_ and spares SVs for subsequent release. By contrast, we find that for inter-train intervals of 1-3 s, the PPR decreased while s_1_ also decreased (**Fig. 2A**). This observation suggests that the response reductions observed with short inter-train intervals are not due to a decrease in the release probability of docked SVs, but to some other process, presumably due to changes in various vesicular pools.

**Figure 2:**
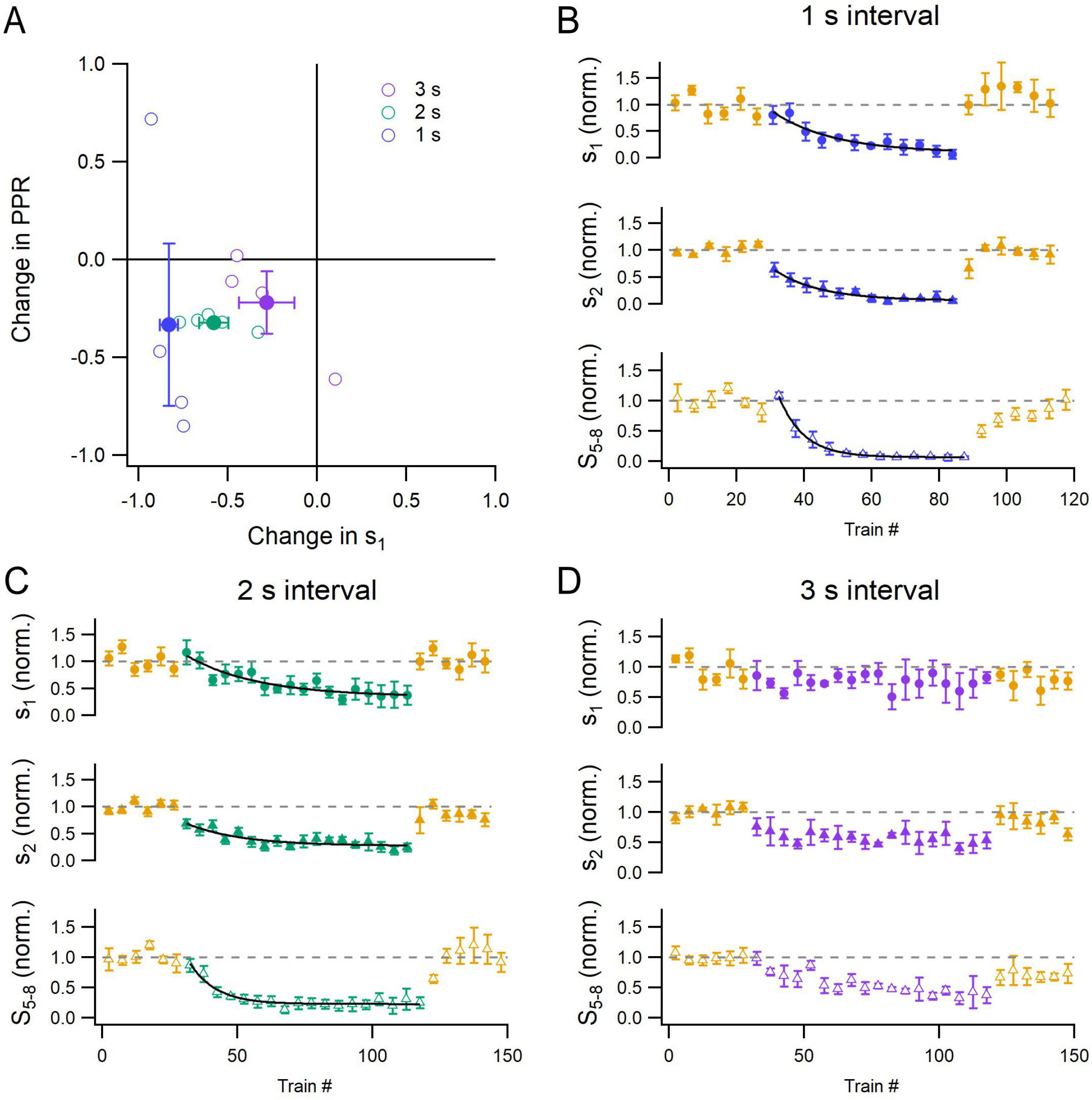
Differential evolution of s_1_, s_2_ and S_5-8_ during changes in the inter-train interval. **A**: Changes in PPR during steady-state release with short inter-train intervals, plotted against the corresponding changes in s_1_. Open circles represent individual experiments. Closed circles indicate the average values for each time interval. **B-D**: Differential evolution of s_1_, s_2_, and S_5-8_ when the inter-train interval was reduced to 1 s (B), 2 s (C), and 3 s (D). Each data point indicates the average of 5 consecutive trains. Traces were fitted with exponential functions (black), yielding time constants of 19 trains, 14 trains, and 7 trains for s_1_, s_2_, and S_5-8_ for 1 s interval, and of 28 trains, 21 trains, and 10 trains for s_1_, s_2_, and S_5-8_ for 2 s interval. n = 4-5 for each time interval. Error bars show +/-SEM.

Previous work has shown that under our recording conditions (3 mM external calcium and 1 mM TEA), the release probability of docked SVs is close to 1 (>0.9: Malagon et al., 2020). The number of SVs released by the first AP in a train, s1, is then an indicator of the size of the docked SV pool, while the number of SVs released by the second AP, s2, is an indicator of the size of the replacement pool. Finally, the cumulative number of SVs released by the 5^th^-8^th^ APs, S_5-8_, is an indicator of the size of a small pool of SVs placed between the recycling pool and the replacement pool, called the intermediate pool (IP; **Fig. 1A**; Tran et al., 2022). Altogether, it is possible to use synaptic response to 8-AP trains to evaluate the evolution of the various SV pools outlined in **Fig. 1A**.

Group results of analysis exploring changes in pool sizes as a function of the inter-train interval are shown in **Fig. 2B-D** (each data point is the average of 5 consecutive trains; n = 4-5 for each interval). For both 1 and 2 s time intervals, the extent of reduction was smaller for s1 than for s2 (by 58 ± 8% vs. 72 ± 5 %, respectively, for 2 s interval; ppt = 0.010; **Suppl. Fig. 1C**), and this smaller reduction was associated with a slower time course (**Fig. 2B-C**: compare the two top panels in each figure). The reduction in s1 was also more moderate than for S5-8 (**Fig. 2B-C**: compare the top and bottom panels; **Suppl. Fig. 1D**). Similar differential reductions of s1 compared to s2 and S5-8 were observed with the 3 s inter-train interval, although in this case, s1 was not significantly reduced (2.2 ± 0.8 vesicles at baseline vs. 1.5 ± 0.4 vesicles during steady state of 3 s interval; ppt = 0.2; **Fig. 2D and Suppl. Fig. 1C-D**). A general rule that emerged was that, during the decrease of synaptic response at short inter-train intervals, the reduction in s1 was smaller than that of s2 and of S5-8. In view of our previous work (Tran et al., 2022), this result indicates that the synapse was likely to be less depleted of docked SVs than of either replacement or IP SVs.

### Activation of a rapid vesicular recycling route with short inter-train intervals

Previous work at hippocampal synapses and neuromuscular junctions suggests that during high frequency stimulation, a rapid route of vesicle recycling is activated to help maintain effective transmission (Ertunc et al., 2007; Maeno-Hikichi et al., 2011; Pyle et al., 2000; Richards et al., 2003; Sara et al., 2002). This route can be blocked by the potent V-ATPase inhibitor folimycin (Ertunc et al., 2007; Maeno-Hikichi et al., 2011), which inhibits the reacidification of recently endocytosed SVs and thus prevents them from being filled with neurotransmitter. In the next set of experiments, we tested whether a rapid route of vesicle recycling is also activated at GC-MLI synapses during short inter-train intervals.

After obtaining a series of baseline responses with 10 s intervals, we bath-applied folimycin (80-160 nM; K_d_ = 0.02 nM; Droese et al., 1993) for 20 min, without stimulation. We then resumed stimulation in folimycin with 2 s inter-train intervals. As illustrated in the experiment of **Fig. 3A**, the first responses in folimycin (green traces) were similar to the baseline (yellow traces), but subsequent responses were quickly inhibited. In this example, the reduction of total SV release per train during stimulation with 2 s intervals had a time constant of only 10 trains (or 20 s; **Fig. 3B**). When the inter-train interval was returned to 10 s, partial recovery to the baseline level was observed (orange; **Fig. 3A-B**).

**Figure 3:**
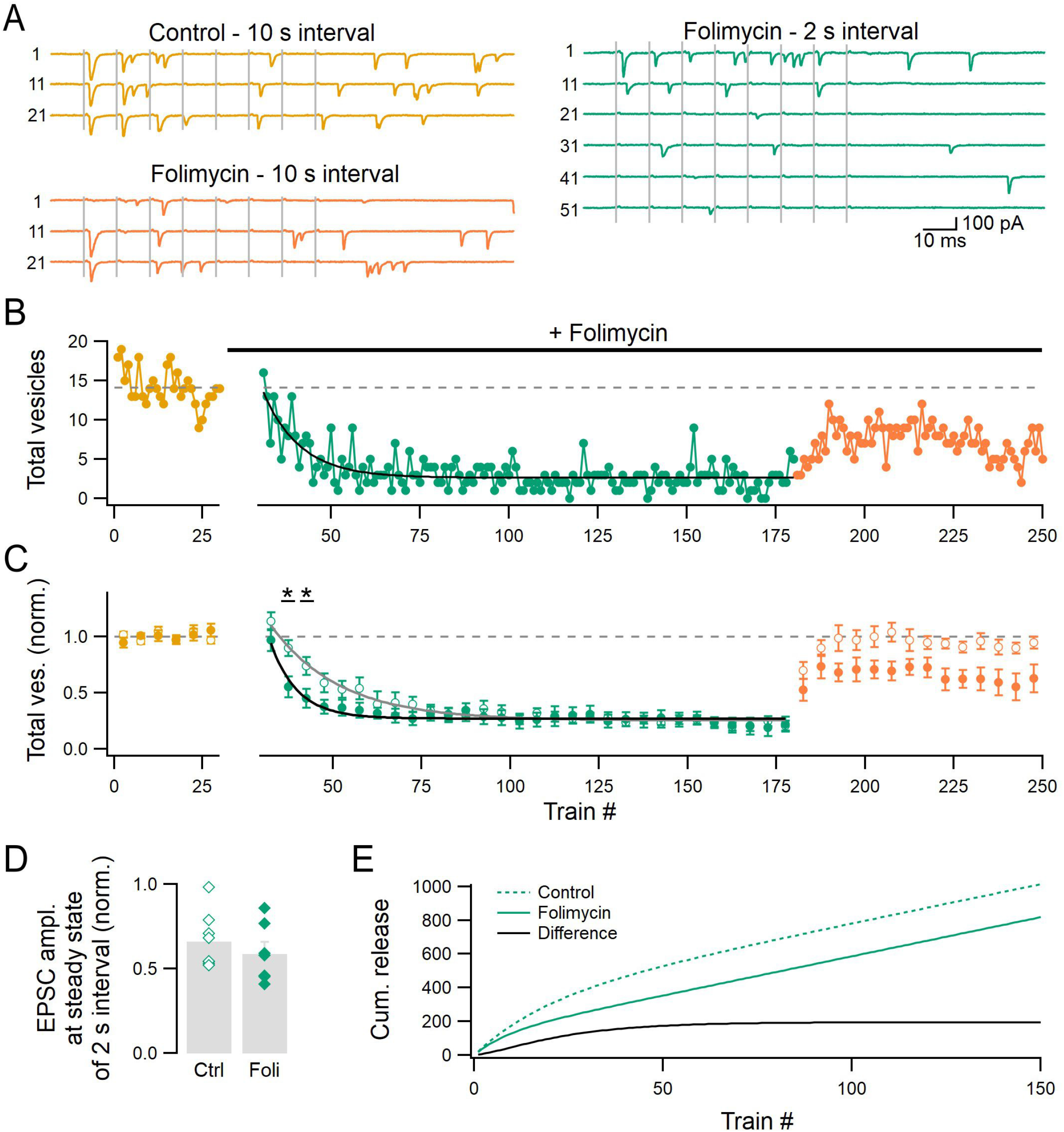
Folimycin accelerates the reduction of release during short inter-train intervals. **A**: An example recording. Postsynaptic responses to 8-AP trains (100 Hz) were first recorded with a 10 s inter-train interval (yellow). Folimycin was then added to the bath and the recording was resumed with an inter-train interval of 2 s (green). Once 150 trains had been obtained with 2 s intervals, the inter-train interval was returned to 10 s (orange). The train number relative to each recording condition is indicated. **B**: The total number of SVs released per train, plotted against the train number for the same experiment. The reduction of release during 2 s intervals was fitted with an exponential function (black), with a time constant of 10 trains. **C**: Group data comparing the evolution of cumulative SV release in control experiments (with or without DMSO; open circles; n = 9) and in experiments with folimycin (closed circles; n = 7). Each data point indicates the average of 5 consecutive trains. Grey and black curves represent exponential fits to the control and folimycin data during the stimulation period with 2 s intervals. The corresponding time constants are 19 and 8 trains. **D**: The average amplitude of EPSCs during steady-state release with 2 s intervals in control vs. folimycin experiments. Values had been normalized with respect to baseline. **E**: The integrated numbers of released SVs as a function of train number during the stimulation period with 2 s intervals in control vs. folimycin experiments, and the difference between them. Green traces were obtained by integrating the exponential fits to the data in C.

Group results show no significant difference in the total number of SVs released per train between control and folimycin experiments at the beginning of the stimulation period with 2 s intervals (train numbers 1-5; 1.1 ± 0.1 vs. 1.0 ± 0.1 of baseline; p-value for unpaired t-test pt = 0.2; n = 7 for folimycin and 9 for control experiments; **Fig. 3C**; each data point is the average of 5 consecutive trains; closed circles indicate SV release in experiments with folimycin; open circles correspond to control experiments with or without DMSO; see **Methods** and **Suppl. Fig. 2E-G**). The steady-state levels of total SV release per train were also not different between control and folimycin experiments (0.26 ± 0.03 vs. 0.26 ± 0.06 of baseline, respectively; pt = 0.9). However, between train numbers 6 and 15 with 2 s inter-train intervals, the reduction in SV release was significantly larger in the presence of folimycin (p_t_ ≤ 0.02). Consistently, the response decayed with a time constant of 8 trains (or 16 s) in folimycin, compared to 19 trains (or 38 s) in control experiments. These results indicate that folimycin inhibits a route of vesicle supply that is mostly active at the onset of the stimulation period with 2 s intervals. It should be noted here that while the mean quantal amplitude was reduced with 2 s inter-train intervals (to be described in a separate publication), this reduction was not augmented by folimycin (by 34 ± 5% in control vs. 41 ± 7% in folimycin; pt = 0.4; **Fig. 3D**). Therefore, it was unlikely that partially filled SVs were present and released during the steady state of 2 s intervals. The majority of SVs that were released during this state likely came from an upstream pool and had been prefilled with glutamate before addition of folimycin.

**Fig. 3E** displays the integrated numbers of released SVs during the entire stimulation period with 2 s intervals for control and folimycin experiments, and the difference between them. As suggested earlier (Hori and Takahashi, 2012; Ikeda and Bekkers, 2009; Qiu et al., 2015; Wojcik et al., 2004; Zhou et al., 2000), SVs devoid of glutamate can still undergo exocytosis. Therefore, SVs are likely released in equal numbers under control conditions and in folimycin. However, in the presence of folimycin, recently endocytosed SVs cannot produce any signal upon re-release as they lack glutamate. Accordingly, the difference in SV counts during the stimulation period with 2 s intervals between control and folimycin experiments represents a rapid route of vesicle recycling (Ertunc et al., 2007). This route is activated upon the onset of 2 s inter-train intervals and it allows the release of supplementary SVs after a lag of about 10 s (5 trains). This indicates that the first SVs that are released by the rapid route have undergone endocytosis ≤10 s before release. The fact that the filling of SVs with glutamate has a time constant of ~10 s (Hori and Takahashi, 2012) explains the strong sensitivity of this recycling route to folimycin. As seen in **Fig. 3E** (black curve), the difference in the integral of SV counts with 2 s intervals between control and folimycin experiments reaches a plateau after ~70 trains, suggesting that this route has a limited capacity and can only provide ~200 SVs during stimulation with 2 s intervals. During the steady-state release with 2 s intervals, the activity of this route is minimal, although unlikely to be zero (see below).

Upon the return of the inter-train interval to 10 s, the total number of SVs released per train recovered to the baseline level in control experiments (**Fig. 3C**). The response in folimycin followed a similar return to baseline, but it deviated from the control after ~10 trains and started to decrease gradually. This likely reflects the presence of a significant proportion of empty SVs in vesicular pools responsible for RRP replenishment at this late stage of the experiments.

### Folimycin abolishes the differential depression of s1, s2 and S5-8 with short inter-train intervals

The evolutions of s1, s2 and S5-8 in control and folimycin experiments are compared in **Fig. 4A-C** (each data point is the average of 5 consecutive trains). The total duration of the stimulation period with 2 s inter-train intervals was 5 min here, compared with 3 min in **Fig. 2C**. During this period, as in **Fig. 2C**, the control data displayed a more moderate decrease for s_1_ than for s_2_ and S_5-8_ (0.56 ± 0.12, 0.21 ± 0.03, and 0.15 ± 0.04 of baseline, respectively; p_pt_ < 0.01; open green symbols in **Fig. 4A-C** and in **Fig 4D**). When returning to the 10 s inter-train interval, s_1_ recovered rapidly and displayed a significant enhancement during the first 5 min (to 150 ± 20% of the baseline value; *p*_pt_ = 0.03; open orange circles in **Fig. 4A** and in **Fig. 4E**). In comparison, both s_2_ and S_5-8_ recovered more slowly and neither of them displayed an overshoot (s2 recovered to 99% of the initial value after ~1 min and S_5-8_ recovered to 88% of the initial value after >5 min). These results indicate a relatively long-lasting enhancement of s_1_ in control conditions after extended stimulation with 2 s inter-train intervals. This number is the product of the number of docking sites, N, of the probability of occupancy of docking sites, δ, and of the probability of release per docked SV, p_r_ (Scheuss and Neher, 2001). In our recording conditions, p_r_ was very close to 1 (Malagon et al., 2016).

**Figure 4:**
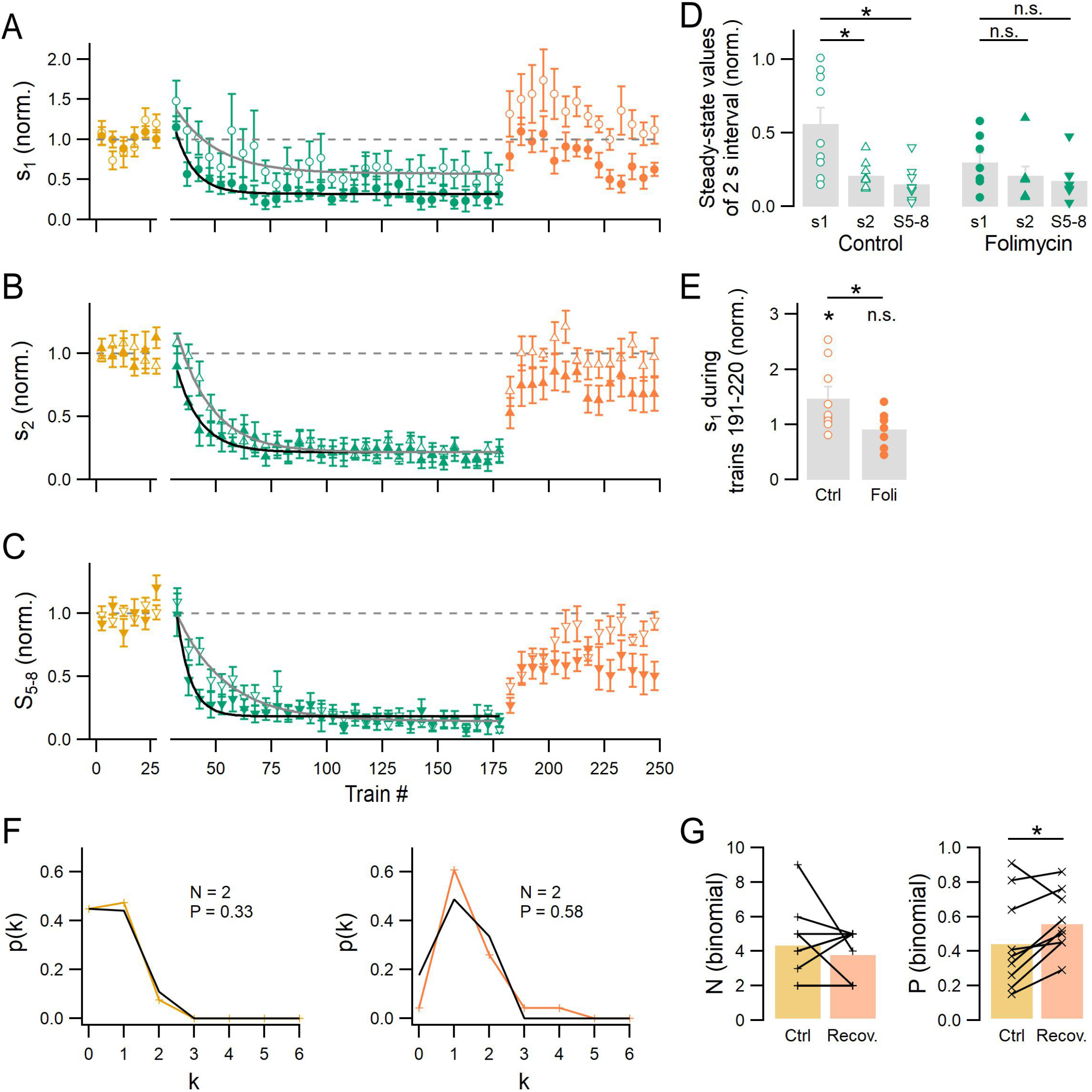
Folimycin abolishes the differential changes between s1 and either s2 or S5-8 during short inter-train intervals. **A-C**: Group data comparing the evolution of s1 (A), s2 (B), and S5-8 (C) in control experiments (with or without DMSO; open symbols; n = 9) vs. in experiments with folimycin (closed symbols; n = 7). In each experiment, responses were first obtained during a baseline period with 10 s inter-train intervals (yellow), then during a test period with 2 s intervals (green), and finally after the return of the inter-train interval to 10 s (orange). Each data point indicates the average value of 5 consecutive trains. Grey curves represent exponential fits to control data, with time constants of 19 trains for s1, 16 trains for s2, and 22 trains for S5-8. Black curves represent exponential fits to folimycin data, with time constants of 8 trains for s1, 11 trains for s2, and 6 trains for S5-8. **D**: The steady-state values of s1, s2, and S5-8 during 2 s intervals in control and in folimycin experiments. **E**: The value of s1 when the inter-train interval was returned to 10 s in control vs. folimycin experiments. **F:** An example of the probability distribution p(k) to observe k vesicular release events after the first AP during baseline recording (yellow; left) and during trains 191-220 (orange; right). The number of docking sites (N) and the release probability per docking site (P = δ pr) obtained from binomial fits (black) are indicated. **G:** The values of N (left) and P (right) during the baseline recording and after the inter-train interval was returned to 10 s.

Binomial analysis of the probability distribution of release after the first AP showed no significant change in N during the period of s1 potentiation (p_pt_ = 0.5; **Fig. 4F-G**; see also Böhme et al., 2019). By contrast, the probability of release per docking site was increased by 50 ± 20% (from 0.45 ± 0.10 to 0.57 ± 0.06; ppt = 0.048; **Fig. 4F-G**). Altogether, our data suggest an increase in δ. The value of δ has been estimated to be 0.47 in 3 mM external calcium concentration (Malagon et al., 2020). Therefore, the increased δ value at the end of the 5 min stimulation period with 2 s inter-train intervals is estimated to be 1.5 × 0.47 = 0.71. Enhanced docking was observed during a period of 6-7 minutes after returning to the 10 s inter-train interval, before subsiding near the end of the recordings (**Fig. 4A**). No such excess was apparent in the data of **Fig. 2C**, indicating that the protracted increase in δ illustrated in the control data of **Fig. 4E** requires a period of >3 min of stimulation with 2 s inter-train intervals.

The time constants of s_1_, s_2_, and S_5-8_ depression during the stimulation period with 2 s inter-we tested the effectstrain intervals were consistently shorter in folimycin (respectively 8, 11 and 6 trains; closed green symbols in **Fig. 4A-C**) than in control (respectively 19, 16 and 22 trains; open green symbols in **Fig. 4A-C**). This suggests that rapidly recycled SVs are returned to a pool upstream of the RRP (the IP or recycling pool; see below). Nevertheless, folimycin had differential effects on various SV pools, as it abolished the differential reduction of s_1_ compared to that of s_2_ and of S_5-8_ (s_1_ = 0.30 ± 0.08, s_2_ = 0.21 ± 0.07, and S_5-8_ = 0.17 ± 0.06 of baseline; p_pt_ > 0.1; closed symbols in **Fig. 4D**). When the inter-train interval was returned to 10 s, no enhancement was observed for s1 in folimycin (s_1_ = 0.9 ± 0.1 of baseline; p_pt_ = 0.5; closed orange circles in **Fig. 4A** and **Fig. 4E**). As s_1_ resulted from the release of docked SV_s_, the majority of which underwent exocytosis after the first AP in our recording conditions, we propose that after entering the RRP, rapidly recycled SVs are preferentially docked.

As a control for the mechanism of action of folimycin, we tested the effects of Rose Bengal, a potent membrane-permeable inhibitor of vesicular glutamate uptake (IC_50_ = 19 nM; Ogita et al., 2001). As Rose Bengal has only a small effect on the membrane potential generated by V-ATPase at the concentration used (1 μM; IC_50_ = 4.3 μM; Ogita et al., 2001), its action should be largely independent of that of folimycin. As illustrated in **Suppl. Fig. 3**, bath-application of Rose Bengal mimicked in all respects the effects of folimycin incubation, validating our interpretation of folimycin results.

### MLCK activation underlies rapid vesicle recycling

At the neuromuscular junction, activation of the myosin light chain kinase (MLCK) triggers a rapid route of vesicle recycling that is sensitive to folimycin (Maeno-Hikichi et al., 2011; Polo-Parada et al., 2001, 2004, 2005). To investigate whether MLCK is required for the rapid recycling of SVs at GC-MLI synapses, we next examined the effects of the MLCK inhibitor ML-9 (Polo-Parada et al., 2004; Saitoh et al., 1987).

Bath-application of ML-9 (25 μM; K_i_ = 4 μM; Saitoh et al., 1987) for 10 min did not affect SV release during 10 s inter-train intervals (s_1_ = 1.8 ± 0.3 vs. 2.0 ± 0.4 vesicles before vs. after ML-9 addition, respectively; total SV release per train = 17 ± 1 vs. 16 ± 2 vesicles; p_pt_ > 0.1; n = 5; compare yellow and pink traces in example in **Fig. 5A-B**; group results in **Fig. 5C**). However, significant differences between control and ML-9 experiments appeared when the inter-train interval was reduced to 2 s (green traces in example in **Fig. 5B**). The decay of the total SV release per train was faster in ML-9 than in control (**Fig. 5D**; closed circles indicate SV release in experiments with ML-9; open circles correspond to control experiments with or without DMSO). The time constant of this decay was 8 trains (or 16 s), markedly smaller than that obtained in control (19 trains or 38 s), but identical to that obtained in folimycin (**Fig. 3C**). Similar to folimycin, ML-9 did not affect the steady-state level of total SV release per train during 2 s intervals (0.26 ± 0.03 vs. 0.29 ± 0.04 of baseline in control and in ML-9, respectively; pt = 0.4). Nonetheless, unlike in folimycin, the total number of SVs released per train in ML-9 fully recovered when the inter-train interval was returned to 10 s.

**Figure 5:**
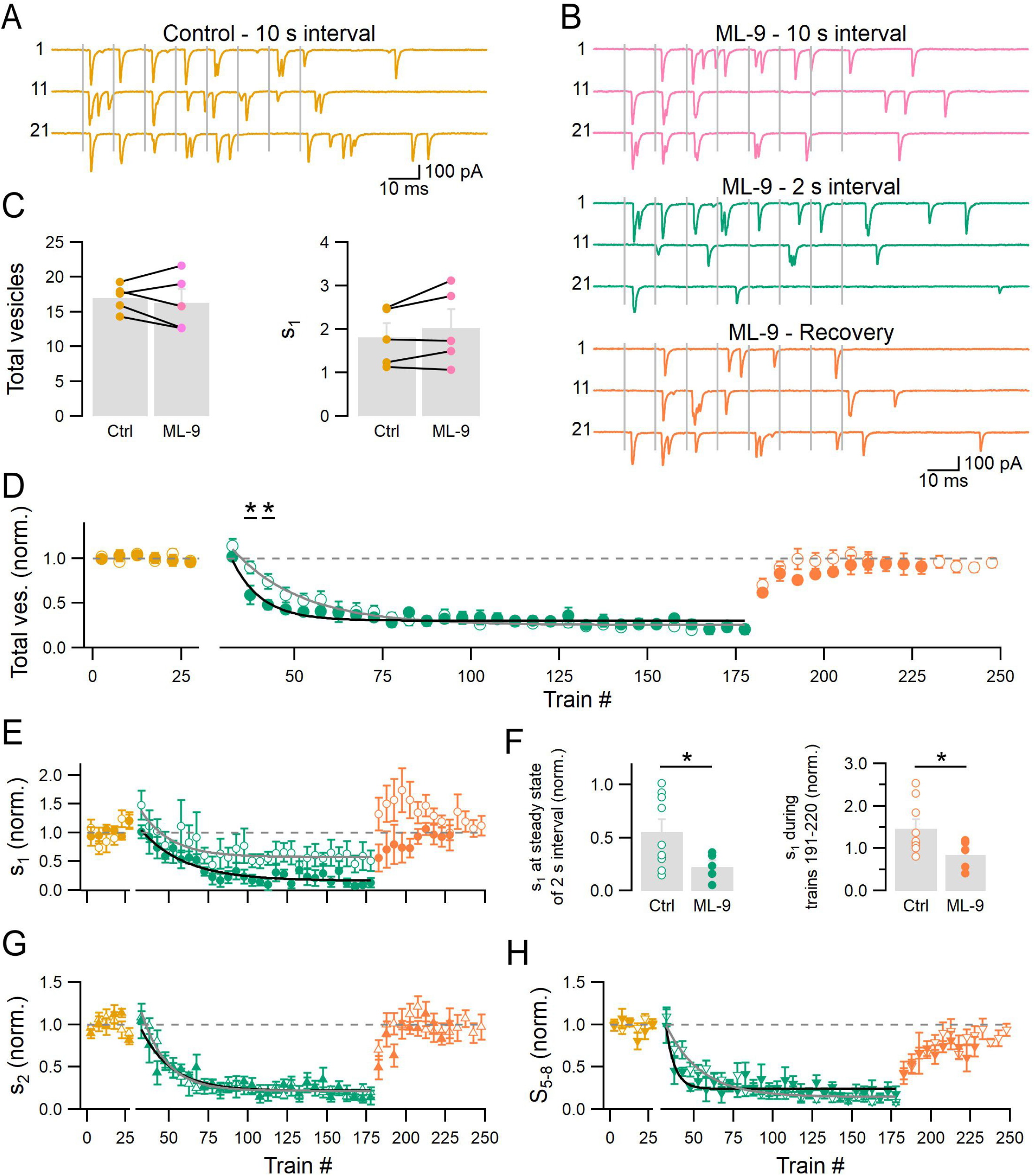
ML-9 accelerates the reduction of release and decreases the steady-state level of s1 during short inter-train intervals. **A-B**: Example recording showing postsynaptic responses to 8-AP trains before (A) and after (B) addition of ML-9 with 10 or 2 s inter-train intervals. **C:** Group results comparing the total number of SVs released per train (left) and s1 (right) before vs. after ML-9 addition. **D**: Group results comparing the evolution of total SV release per train in control experiments (with or without DMSO; open symbols; n = 9) vs. in experiments with ML-9 (closed symbols; n = 5). Each data point indicates the average value of 5 consecutive trains. Data for ML-9 had been normalized to the baseline values before addition of ML-9. Grey and black curves represent exponential fits to the control and ML-9 data, with time constants of 19 trains and 8 trains, respectively. **E**: Similar to D but for s1. The time constants of exponential fits are 19 trains for control and 28 trains for ML-9. **F**: The value of s1 during steady-state release with 2 s intervals (left) and when the inter-train interval was returned to 10 s (right) in control vs. ML-9 experiments. **G**: Similar to D but for s2. The time constants of exponential fits are 16 trains for control and 19 trains for ML-9. **H**: Similar to D but for S5-8. The time constants of exponential fits are 22 trains for control and 5 trains for ML-9.

The steady-state value of s_1_ during 2 s intervals was smaller in ML-9 than in control experiments (0.23 ± 0.06 vs. 0.56 ± 0.12 of baseline, respectively; p_t_ = 0.03; green circles in **Fig. 5E** and **Fig. 5F**, left). Similarly, the potentiation of s_1_ observed during the first 6-7 minutes following the return to 10 s intervals in control conditions (1.5 ± 0.2 of baseline) was abolished by ML-9 (0.8 ± 0.2 of baseline; p_t_ = 0.03; orange circles in **Fig. 5E** and **Fig. 5F**, right). Therefore, like folimycin, ML-9 inhibited the enhanced docking caused by stimulation with short inter-train intervals. ML-9 also accelerated the run-down of S5-8 (time constant = 5 trains in ML-9 vs. 22 trains in control; **Fig. 5H**). In contrast, the evolution of s_2_ was comparable in ML-9 and in control (**Fig. 5G**). Altogether, these results suggest that in GC-MLI synapses, as in the neuromuscular junction, activation of MLCK underlies the rapid recycling of SVs during enhanced activity. They also substantiate the notion that docking is prioritized by this route of vesicle recycling.

### Effects of blocking dynamin-dependent endocytosis

Since inhibition of V-ATPase with folimycin did not affect the steady-state level of total SV release per train during 2 s intervals, it remained unclear what mechanisms underlie this state of release. To investigate the importance of dynamin-dependent endocytosis during the rapid recycling of SVs and during the steady-state release with short inter-train intervals, we examined the effects of the potent dynamin blocker dyngo-4a (IC_50_ = 0.4 μM; McCluskey et al., 2013).

Bath-application of dyngo-4a (30 μM) for 20 min did not have a strong effect on SV release during stimulation with 10 s inter-train intervals (**Suppl. Fig. 4**; n = 5). However, in a separate set of experiments where the inter-train interval was reduced to 2 s after addition of dyngo-4a, the total number of SVs released per train ran down with a time constant of 9 trains (or 18 s; closed green circles in **Fig. 6A**). This value is similar to that obtained with folimycin or ML-9 (8 trains), and smaller than that obtained in control experiments (19 trains). In the presence of dyngo-4a, the steady-state level of total SV release per train during 2 s intervals was also lower than that in control (0.07 ± 0.06 vs. 0.25 ± 0.03 of baseline, respectively; pt = 0.001; **Fig. 6A**). Additionally, the total SV release per train recovered only weakly when the inter-train interval was returned to 10 s (closed orange circles in **Fig. 6A**).

**Figure 6:**
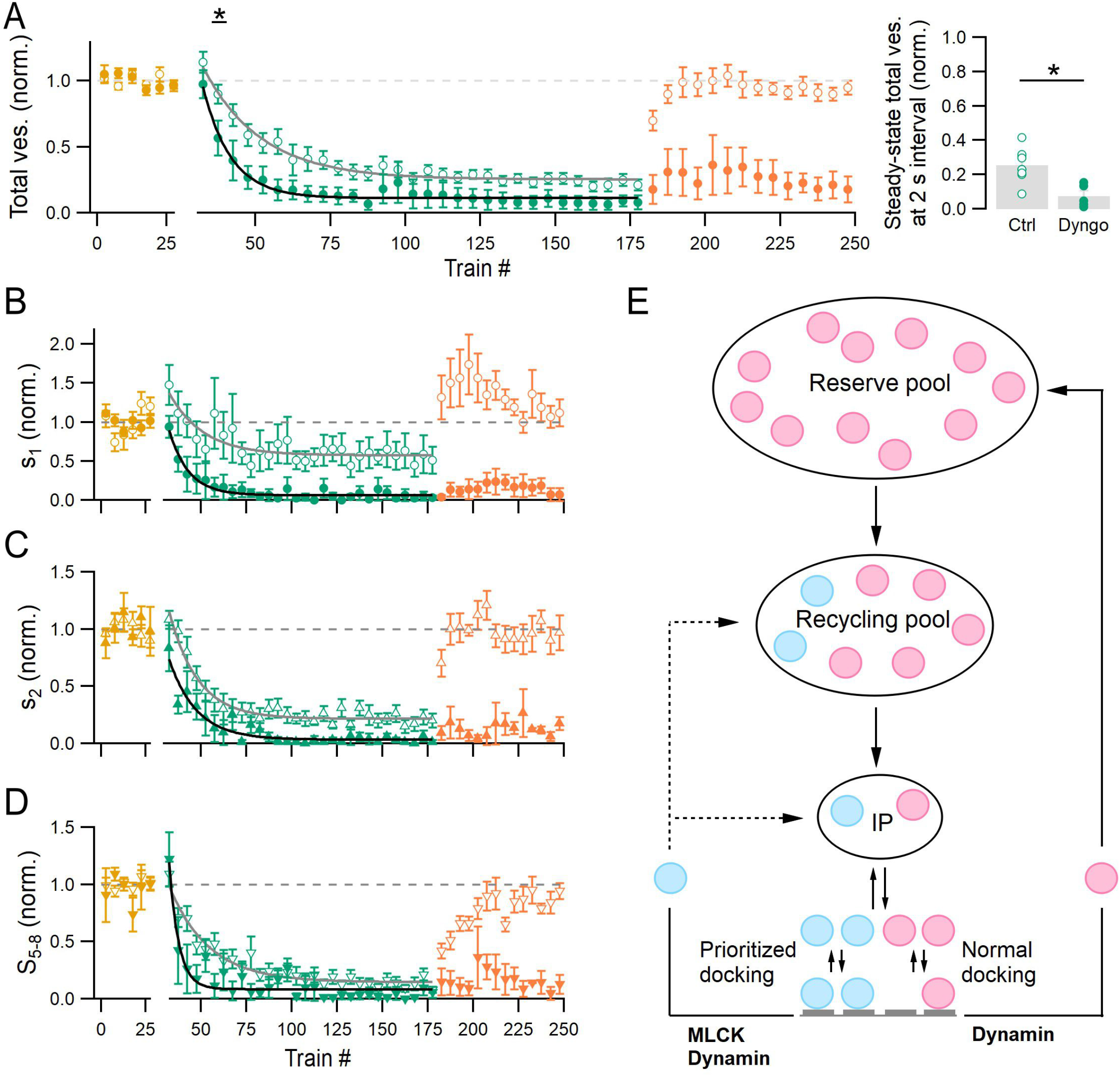
Dyngo-4a accelerates the reduction of release during short inter-train intervals. **A, left**: The evolution of total SV release per train in control experiments (open symbols; n = 9) vs. in experiments with dyngo-4a (closed symbols; n = 5). In each experiment, responses were first obtained with a 10 s inter-train interval (yellow), then with a test interval of 2 s (green), and finally with a 10 s interval (orange). Addition of DMSO and dyngo-4a was done after the baseline recording. Each data point indicates the average value of 5 consecutive trains. Grey and black curves represent exponential fits to the control and dyngo-4a data, with time constants of 19 and 9 trains, respectively. **A, right**: The steady-state value of total SV release per train during 2 s intervals in control vs. dyngo-4a. **B:** Similar to A but for s1. Exponential fits had time constants of 19 trains for control and 11 trains for dyngo-4a. **C:** Similar to A but for s2. The time constant of exponential fits are 16 and 15 trains, respectively.**D:** Similar to A but for S5-8. The time constants of exponential fits are 22 and 5 trains, respectively. **E**: Schematic of two SV recycling pathways in GC-MLI synapses. The central column represents a model of SV replenishment comprising a reserve pool of infinite size, a recycling pool that can be depleted during stimulation with short inter-train intervals, an intermediate pool (IP), and the RRP (including replacement SVs, above, and docked SVs, below; 4 docking sites are depicted, with a total of 7 SVs/AZ). The right loop represents a recycling pathway that does not require SVs to be rapidly refilled with neurotransmitter and thus displays a very weak sensitivity to folimycin. This pathway operates under standard conditions as well as during stimulation with shorter intervals. It presumably returns SVs to the reserve pool. The left loop represents a recycling pathway that requires SVs to be quickly refilled so that they can be re-released soon after endocytosis. This pathway is thus highly sensitive to folimycin. It is activated during enhanced activity (through activation of MLCK), and returns SVs to the recycling pool and/or the IP. SVs that are recycled through this pathway have an increased probability of docking (see text). Both recycling pathways require dynamin-dependent endocytosis.

**Fig. 6B-D** compares the evolution of s_1_, s_2_, and S_5-8_ in dyngo-4a vs. control experiments. Dyngo-4a significantly reduced the steady-state levels of s_1_, s_2_, and S_5-8_ during 2 s inter-train intervals (in dyngo-4a and in control respectively: s_1_ = 0.06 ± 0.05 vs. 0.56 ± 0.12 of baseline; s_2_ = 0.04 ± 0.02 vs. 0.21 ± 0.03 of baseline; S_5-8_ = 0.05 ± 0.03 vs. 0.15 ± 0.04 of baseline; pt ≤ 0.04). Moreover, s_1_, s_2_, and S_5-8_ all failed to recover upon the return of the inter-train interval to 10 s.

The similar decay time courses of the total SV release per train during stimulation with 2 s inter-train intervals in dyngo-4a, folimycin, and ML-9 suggest that the rapid recycling of SVs requires activation of MLCK and efficient functioning of dynamin and V-ATPase. In addition, the strong, largely irreversible inhibition of release observed in dyngo-4a suggests that efficient dynamin-dependent endocytosis is necessary to maintain synaptic transmission during prolonged stimulation with short inter-train intervals, by providing an adequate supply of SVs (Ferguson et al., 2007; Newton et al., 2006), and/or enabling release site clearance for subsequent exocytosis (Hosoi et al., 2009; Hua et al., 2013; Kawasaki et al., 2000; Mahapatra et al., 2016; Wen et al., 2018; Wu et al., 2009).

## Discussion

### Main findings

In the present work, we show that the responsiveness of GC-MLI synapses to successive AP trains can be modulated on a minute timescale by changing the inter-train interval. This modulation is mainly driven by changes in the size of the recycling pool of SVs, with a resting value that can be estimated at 180 SVs per AZ (see **Supplementary Information**). When the inter-train interval is lowered to ≤3 s, the size of the recycling pool decreases gradually, but this reduction is counteracted by activation of a rapid recycling pathway. This pathway is strongly sensitive to folimycin and dyngo-4a. It requires activation of MLCK and generates ~200 SVs per AZ during stimulation with 2 s inter-train intervals. Importantly, rapidly recycled SVs display an enhanced probability of docking, compared to pre-existing SVs. Such prioritized docking can last for minutes after prolonged stimulation with short inter-train intervals.

### Gradual depletion of recycling pool upon repeated AP train stimulation

During sensory stimulation, cerebellar granule cells fire short but high-frequency trains of APs (Chadderton et al., 2004). In the present work, we show that the responsiveness of GC-MLI synapses to such AP trains remains constant for inter-train intervals of ≥10 s. However, for inter-train intervals of ≤3 s, the synapse’s responsiveness gradually decreases until it reaches a steady state after a few minutes of repeated stimulation. We propose that the reduction in synaptic output is mainly caused by a depletion of recycling SVs. Upon returning to the standard 10 s inter-train interval, the synapse’s responsiveness recovers, as the recycling pool returns to its resting size. The steady-state level of release during short inter-train intervals presumably reflects the rate of vesicle recruitment from the reserve pool (see below).

We cannot exclude the alternative hypothesis that, during repeated train stimulation, the release of SVs would be hampered by the accumulation of vesicular membrane and/or proteins at the AZ, rather than by the depletion of upstream vesicular pools. Such a scenario would predict a decrease in p, the release probability of docked SVs, following repeated stimulation at 2 s intervals. However, **Suppl. Fig. 5** shows that repeated stimulation at 2 s intervals resulted in a roughly scaled-down si curve, with abolition of facilitation after the second AP, both in control conditions and in folimycin; in contrast, a decreased p scenario would predict a marked change in the shape of the si curve and an emergence of facilitation during the AP train. Moreover, in the presence of dyngo-4a, the reduction of total SV release per train has the same time constant as that in experiments with folimycin or ML-9. If release site clearance had been the main factor underlying the reduction in synaptic output, blockade of all dynamin-dependent endocytic pathways by dyngo-4a should have had a stronger effect on the rate of reduction compared to blockade of only a rapid recycling pathway by folimycin or ML-9. Altogether, the most parsimonious interpretation of our data remains that the reduced synaptic output during repeated stimulation with short inter-train intervals results from recycling pool depletion.

### Two routes of SV supply during stimulation with short inter-train intervals

During stimulation with 2 s inter-train intervals, folimycin fails to alter the initial and the steady-state responses, but it accelerates the transition to the low steady-state level (**Fig. 3**). Therefore, in agreement with previous studies (Ertunc et al., 2007; Maeno-Hikichi et al., 2011; Pyle et al., 2000; Richards et al., 2003; Sara et al., 2002), we propose that there exist two routes of vesicle supply, with one relying on recently endocytosed SVs while the other utilizing prefilled SVs from an upstream pool (**Fig. 6E**). The former route of SV supply is not active during stimulation with 10 s intervals, but it becomes activated during stimulation with 2 s intervals, presumably by presynaptic calcium rises. We find a delay of 10 s between stimulation onset and the time at which folimycin has a significant effect. This delay is consistent with previous studies demonstrating that a full SV cycle from endocytosis to docking takes ≤10 s (Watanabe et al., 2014) and that the reacidification and refilling of SVs with glutamate have time constants of ~5-10 s (Atluri and Ryan, 2006; Granseth et al., 2006; Hori and Takahashi, 2012). The finding that not only s_1_ and s_2_, but also S_5-8_, are sensitive to folimycin (**Fig. 4**) suggests that rapidly recycled SVs are not returned directly to the RRP, but rather to a pool upstream of the RRP, either the IP or the recycling pool (left loop in **Fig. 6E**). Two conclusions can be reached based on the finding that folimycin does not significantly affect the steady-state response during stimulation with 2 s intervals. First, the supply of SVs by the rapid recycling route is limited. More specifically, this route can only generate ~200 SVs/AZ during stimulation with 2 s intervals. However, additional results discussed below indicate that this pathway does not disappear entirely at steady state, but that it then operates at a low rate to maintain a relatively high number of docked SVs.

The second conclusion is that the synapse mainly uses prefilled SVs from an upstream pool during steady-state release with 2 s inter-train intervals. This pool is likely to be replenished by a more distal recycling route (right loop in **Fig. 6E**) because in the presence of folimycin, the loss of glutamate from its SVs only becomes apparent at the very end of our recordings. As its size appears to be larger than that of the recycling pool, we propose that it corresponds to the reserve pool.

### Prioritized docking

We find that during repeated train stimulation with 2 s intervals, s1 decreases less than later responses in a train (s_2_-s_8_; **Fig. 2**). This indicates an increased likelihood of SVs within the RRP to be docked (prioritized docking). As the transition of SVs from a replacement site to a docking site is thought to be reversible (Kusick et al., 2020; Miki et al., 2018; Tran et al., 2022) (**Fig. 6E**), prioritized docking could involve an increased rate constant of docking, a decreased rate constant of undocking, or both. When the inter-train interval returns to 10 s, s1 not only recovers but it also displays an overshoot compared to the baseline value (**Fig. 4**). This enhancement lasts for several minutes after the return to the 10 s interval. It occurs after 150 trains with 2 s intervals, but not after 90 trains with 2 s intervals, suggesting that prioritized docking lasts longer after prolonged stimulation with short intervals.

Strikingly, folimycin abolishes prioritized docking, both during and after stimulation with 2 s intervals (**Fig. 4**). This indicates that prioritized docking occurs exclusively for rapidly recycled SVs (left loop in **Fig. 6E**). Therefore, in the presence of folimycin, newly formed SVs undergo prioritized docking but their exocytosis does not produce any postsynaptic current because they lack neurotransmitter. In contrast, prefilled SVs coming from the reserve pool are docked with a normal probability (right loop in **Fig. 6E**). ML-9 and dyngo-4a, like folimycin, block prioritized docking, but in these cases, the block is due to the absence of rapidly recycled SVs.

### Mechanisms of action of MLCK

#### Endocytosis

The finding that ML-9, a blocker of MLCK, reproduces the effects of folimycin (**Fig. 5**) is in line with extensive results gathered at the neuromuscular junction indicating that MLCK activation is required for the rapid recycling of SVs during high-frequency stimulation (Maeno-Hikichi et al., 2011; Polo-Parada et al., 2001, 2004, 2005). More recently, MLCK blockers have been shown to inhibit endocytosis at the calyx of Held and in hippocampal neurons (Li et al., 2016; Yue and Xu, 2014), suggesting the existence of the same pathway in central synapses. Based on these previous studies, it appears that the rapid recycling of SVs involves several steps including activation of MLCK by calcium/calmodulin, phosphorylation of myosin II, and induction of dynamin-dependent endocytosis.

#### Myosin II-mediated docking

At central synapses, MLCK and myosin II are involved not only in endocytosis, but also in vesicle recruitment to release sites, probably by promoting docking (Hayashida et al., 2015; Lee et al., 2010, 2012; Miki et al., 2016; Ryan, 1999). In light of these earlier studies, our results suggest that following activation by prolonged and repeated calcium elevations, the MLCK pathway exerts two distinct effects on synapses, triggering endocytosis and enhancing SV docking. Since MLCK can be activated by calmodulin, the coexistence of two targets for MLCK is consistent with the suggestion that at the calyx of Held, activation of calmodulin separately enhances endocytosis and vesicle recruitment to the RRP (Yao and Sakaba, 2012). In the present work, we show that while these two effects are distinct, they interact in prioritized docking, since this phenomenon only occurs for recently endocytosed SVs. These results suggest that following MLCK-driven endocytosis, SVs display an identification signal as they enter the RRP, such that only these SVs undergo prioritized docking. The nature of this signal remains at present unknown.

#### Effects of dyngo-4a

As discussed above, the reversible reduction of synaptic output during repeated stimulation with 2 s inter-train intervals likely reflects the reversible depletion of the recycling pool. Therefore, the finding that dyngo-4a prevents the recovery of the synaptic output after stimulation with 2 s intervals indicates that the pool upstream of the recycling pool (i.e., the reserve pool) is compromised by dyngo-4a. This means that dynamin is essential to the distal recycling route which replenishes the reserve pool (right loop in **Fig. 6E**). Due to the longer duration of this pathway, folimycin may not be able to block the refilling of its SVs. This could explain why the effect of folimycin on the steady-state release with 2 s intervals is much weaker than that of dyngo-4a. Additionally, blockade of dynamin-dependent endocytosis by dyngo-4a may prevent the clearance of exocytosed materials from release sites, thereby inhibiting subsequent release (Hosoi et al., 2009; Hua et al., 2013; Kawasaki et al., 2000; Mahapatra et al., 2016; Wen et al., 2018; Wu et al., 2009).

#### Physiological roles of rapid SV recycling and of prioritized docking

The present results suggest that, as the synapse enters a period of short inter-train intervals, the rapid recycling of SVs acts to delay recycling pool depletion and thus preserve the strength of the synaptic response. Furthermore, prioritized docking ensures that, even when vesicular pools are depleted, responses to isolated APs remain close to those obtained in control conditions. Finally, the s1 overshoot observed after prolonged stimulation with 2 s inter-train intervals suggests that prioritized docking could play a role in post-tetanic potentiation, in line with previous findings indicating a post-tetanic enhancement of the fast-releasing SV pool at the calyx of Held (Lee et al., 2010).

## Acknowledgements

We thank Takafumi Miki and Melissa Silva for helpful comments on the manuscript. This work was supported by CNRS (UMR 8118, and UMR 8003), by the European Community (ERC Advanced Grant ‘Single Site’ to A. M., nb. 294509), and by Fondation pour la Recherche Médicale (grant SPF201809007190 to V. T.).

## Author Contributions

V.T. and A.M. designed research; V.T. performed research and analyzed data; and V.T. and A.M. wrote the paper.

## Materials and Methods

### Preparation of acute cerebellar slices

The use and care of experimental animals were in accordance with guidelines of Université de Paris (approval no. D 75-06-07). 200 μm thick sagittal slices were prepared from the cerebellar vermis of Sprague-Dawley rats (P13-17) as described previously(Llano et al., 1991). Slices were incubated at 34 °C for at least 45 min before being used for experiments.

### Recording of a simple GC-MLI synapse

Whole-cell patch-clamp recordings were obtained from molecular layer interneurons (MLIs; comprising both basket and stellate cells). The extracellular solution contained (in mM): 130 NaCl, 2.5 KCl, 26 NaHCO_3_, 1.3 NaH_2_PO_4_, 10 glucose, 3 CaCl_2_, and 1 MgCl_2_. It was equilibrated with 95% O_2_ and 5% CO2 (pH 7.4). The internal recording solution contained (in mM): 144 K-gluconate, 6 KCl, 4.6 MgCl2, 1 EGTA, 0.1 CaCl2, 10 HEPES, 4 ATP-Na, 0.4 GTP-Na (pH 7.3, 300 mosm/l). All recordings were done at 32-34 °C and in the presence of D-APV (50 μM) and gabazine (3 μM) to block NMDA and GABA_A_ receptors, respectively.

A single GC-MLI connection was established as described previously(Miki et al., 2017). Briefly, a glass pipette filled with the high K^+^ internal solution was used to identify a potential presynaptic granule cell. The same pipette was then used for extracellular electrical stimulation of the granule cell, with minimal stimulation voltage. If the resultant EPSCs, particularly those that occurred after a short stimulation train, were homogenous, then it was likely that only one presynaptic cell was stimulated.

Recordings were only accepted as a simple synapse recording after analysis and if the following three criteria were satisfied(Malagon et al., 2016): i) the EPSC amplitude of the second release event in a pair was smaller than that of the first, reflecting activation of a common set of receptors belonging to one postsynaptic density; ii) all the EPSC amplitudes follow a Gaussian distribution with a coefficient of variation < 0.5; and iii) the number of release events during the baseline recording was stable.

Folimycin (Abcam), ML-9 (Tocris), and Dyngo-4a (Tocris) were dissolved in dimethyl sulfoxide (DMSO), and stock solutions were kept at −20 °C. The final DMSO concentration was <0.2 %. Rose Bengal (Tocris) was dissolved in water. The final concentration of folimycin was 80 nM in one experiment, 120 nM in two experiments, and 160 nM in four experiments. The results of these experiments were comparable and thus pooled together in group analysis. Similarly, the results of control experiments in which DMSO was added after baseline recording were not different from those without DMSO addition (**Suppl. Fig. 2E-G**); therefore, they were pooled.

### Decomposition of EPSCs

The time of occurrence and the amplitude of individual release events were determined based on deconvolution analysis, as described previously(Malagon et al., 2016). For each synapse, an mEPSC template was obtained and fitted with a triple-exponential function with five free parameters (rise time, peak amplitude, fast decay time constant, slow decay time constant, and amplitude fraction of slow decay). These five parameters were then used for deconvolution of the template and of individual data traces, producing a narrow spike (called spike template) and sequences of spikes, respectively. Next, each deconvolved trace was fitted with a sum of scaled versions of the spike template, yielding the timing and amplitude of each release event. The amplitude was further corrected for receptor saturation and desensitization, using the exponential relationship between individual amplitudes and the time interval since the preceding release events. Events that were 1.7 times larger than the average mEPSC were split into two(Malagon et al., 2016). The amplitude of each of these two events was chosen to be half of that of the original event, and their timing was separated by 0.1 ms.

s_1_ was determined as the number of SV_s_ released within 5 ms after the 1^st^ AP of a train. s_2_ was the number of SV_s_ released within 5 ms after the 2^nd^ AP. S_5-8_ was the sum of SVs released within 5 ms after the 5^th^-8^th^ APs. The total number of SVs released per train included all release events within 200 ms after stimulation onset.

### Model of recycling pool replenishment

We call R(t) the number of SVs located in the recycling pool at time t, and R_o_ this number under baseline conditions (10 s inter-train intervals). In the presence of folimycin, the recycling of recently endocytosed SVs is muted. The rate of change of R during a stimulation period with a short inter-train interval is then given by:

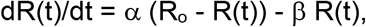

where α and β are rate constants characterizing the exchange of the recycling pool with the reserve pool and with the IP, respectively.

The above equation predicts an exponential decay for R(t) with a time constant of decay

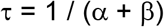

The initial slope of R(t) at time 0 is

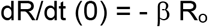

In this equation, β represents the initial loss rate of SVs from the recycling pool when entering the short interval period. ß is inversely proportional to the inter-train interval, so that the above relation between τ and ß is in line with the finding that τ increases linearly as a function of the inter-train interval (**Fig. 1E**; note however that the data in this plot were obtained in the absence of folimycin, therefore they presumably include a component from the folimycin-sensitive pathway which is not considered in the present analysis). Finally, the steady state value of R, R_ss_, is given by

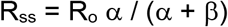

With an inter-train interval of 2 s, the averaged experimental value of τ is 16 s, dR/dt (0) =−8.5 SVs/s, and R_ss_/R_o_ = 0.25. With these values, all parameters of the model can be determined, giving α = 0.0156 s^-1^, β = 0.047 s^-1^, and Ro = 180 SVs.

### Binomial analysis of the probability distribution of release

According to the binomial distribution, the probability that s_1_, the number of SVs released after the first AP, equals k is

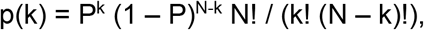

where P is the release probability per docking site and N is the number of docking sites(Malagon et al., 2016). The values of N and P were obtained by minimizing the summed squared deviations between the binomial prediction and data(**Fig. 4F**).

## Notes

### Competing Interest Statement

The authors have declared no competing interest.

