## Supplement File for "Prioritized docking of synaptic vesicles provided by a rapid recycling pathway"

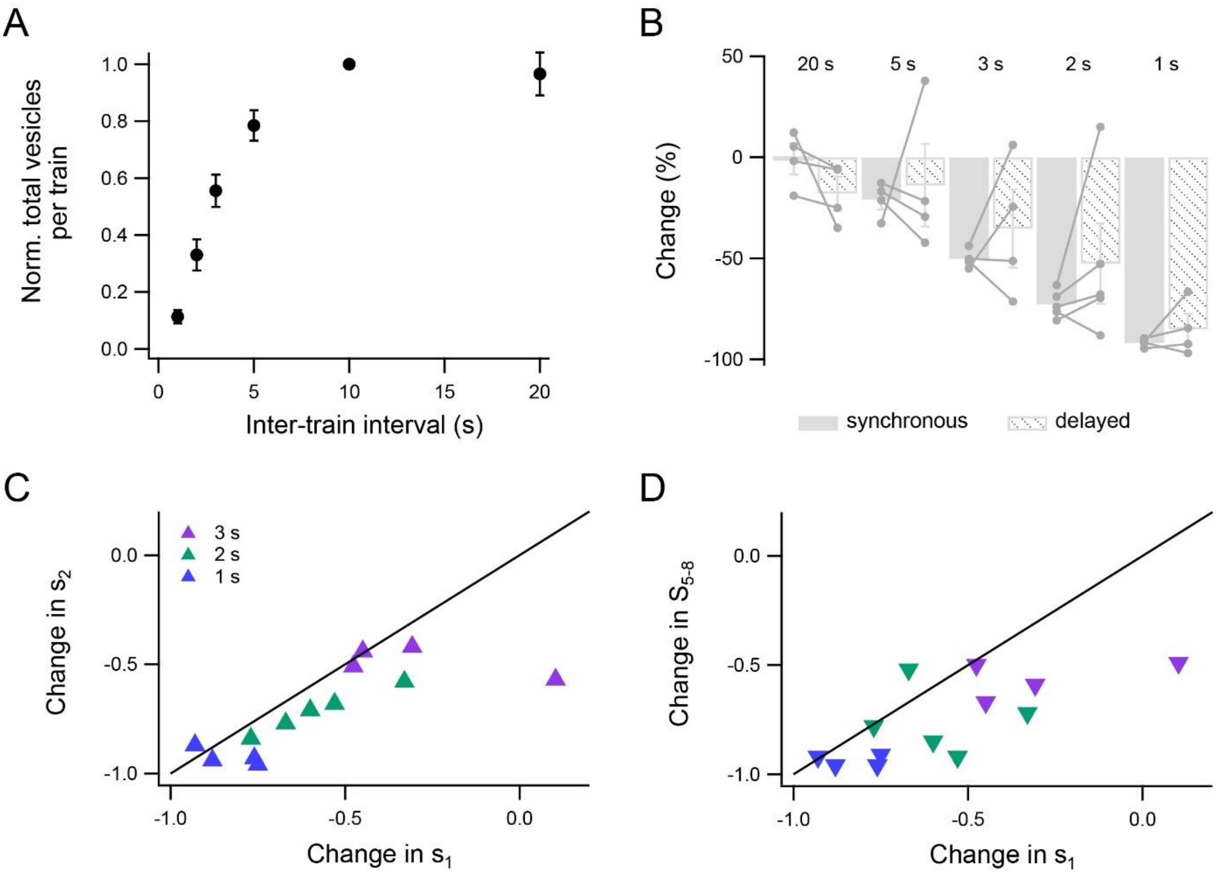


**Supplementary Figure 1: Reduction of SV release during short inter-train intervals**

**A**: The total number of SVs released by an 8-AP train plotted as a function of inter-train interval. In all experiments, the total SV release per train during 10 s inter-train intervals were used to normalize the results obtained with the other inter-train intervals. **B**: Changes in synchronous and delayed release per train at various inter-train intervals, compared to the standard 10 s interval. **C**: Changes in $s_2$ plotted against the corresponding changes in s1. Each data point represents an experiment. The fact that most data points lie below the line of identity (black) indicate that $s_2$ was more reduced than $s_1$ during short inter-train intervals. **D**: Changes in $S_{5\text{-}8}$ plotted against the corresponding changes in $s_1$. Most data points lie below the line of identity (black), suggesting that $S_{5\text{-}8}$ was more reduced than $s_1$.

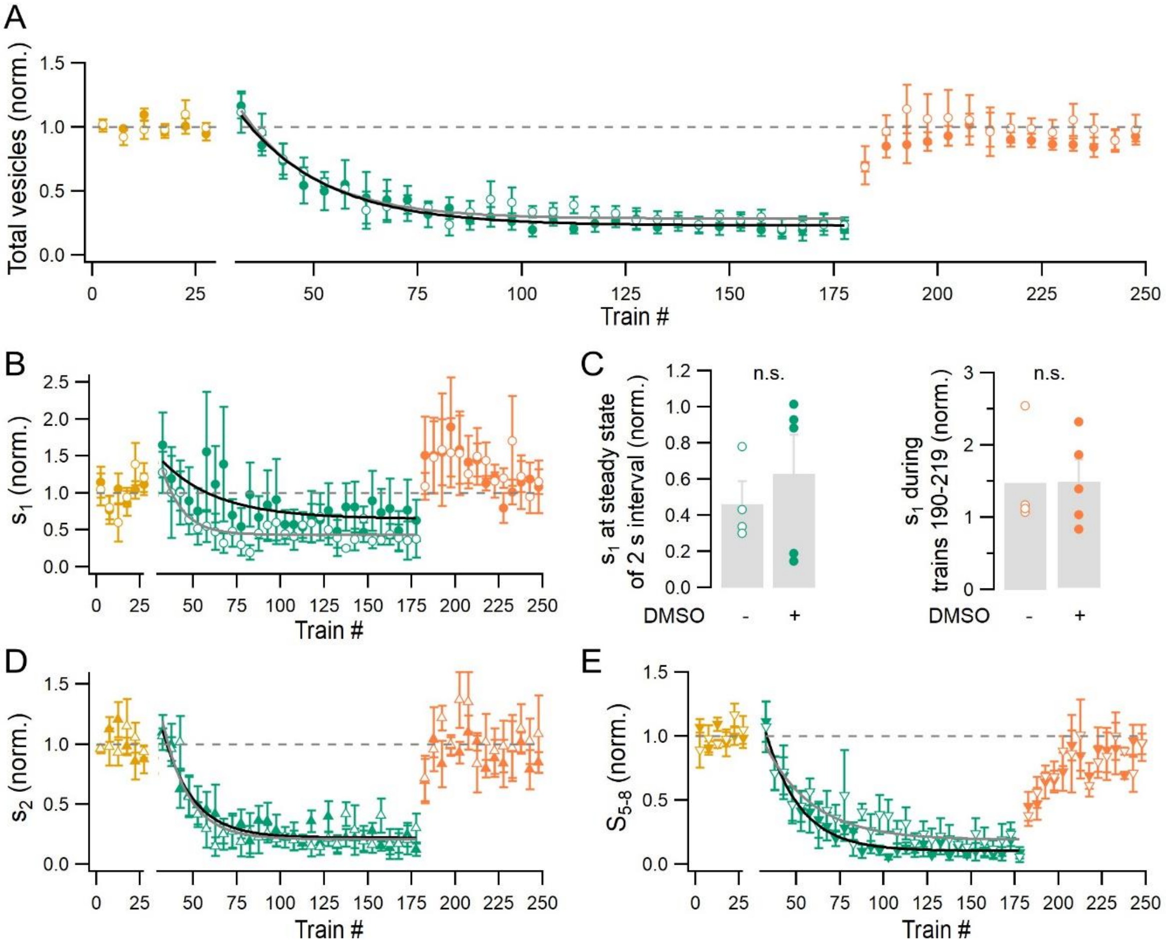


**Supplementary Figure 2: Reduction of SV release during 2 s intervals in control experiments with or without DMSO**

**A**: Group results comparing the evolution of total SV release per train in experiments without DMSO (open symbols; n = 4) vs. in those with DMSO (closed symbols; n = 5). In each experiment, responses were first obtained during a baseline period with 10 s inter-train intervals (yellow), then during a test period with 2 s intervals (green), and finally after the return of the inter-train interval to 10 s (orange). Addition of DMSO was done after the baseline recording had been obtained. Each data point indicates the average value of 5 consecutive trains. Grey and black curves represent exponential fits, with time constants of 18 trains for experiments without DMSO and 20 trains for experiments with DMSO. **B**: Similar to A but for $s_1$. The time constants of exponential fits are 12 trains for experiments without DMSO and 32 trains for those with DMSO. **C**: The value of $s_1$ during steady-state release with 2 s intervals (left) and when the inter-train interval was returned to 10 s (right) in experiments with and without DMSO. **D**: Similar to A but for $s_2$. The time constants of exponential fits are 15 trains for experiments without DMSO and 17 trains for experiments with DMSO. **E**: Similar to A but for $S_{5-8}$. The time constants of exponential fits are 27 trains and 20 trains, respectively.

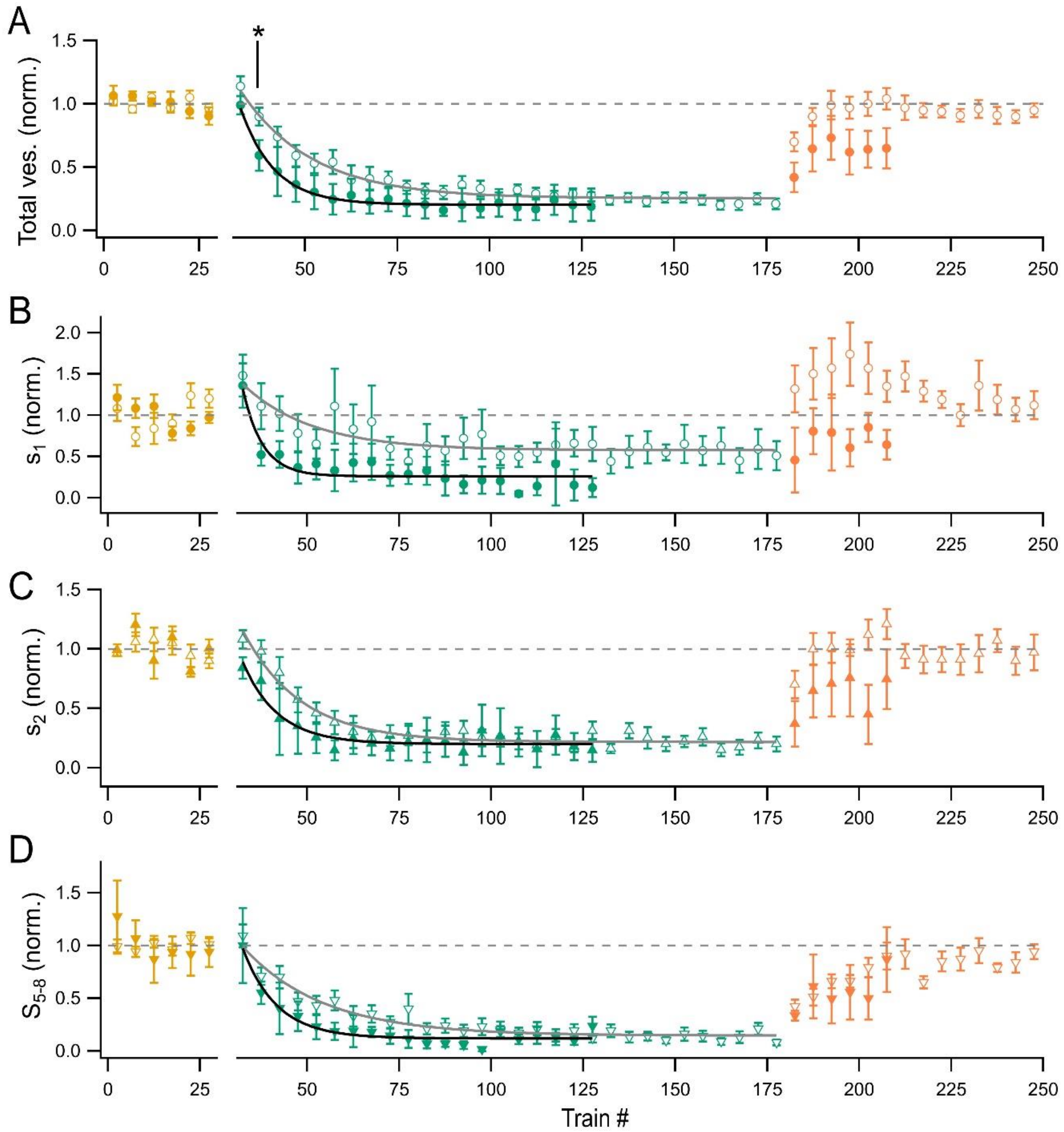


**Supplementary Figure 3: Rose Bengal accelerated the reduction in SV release during 2 s inter-train intervals**

**A**: Group results comparing the evolution of total SV release per train in control experiments (with or without DMSO; open symbols; n = 9) vs. in experiments with Rose Bengal (1 µM; closed symbols; n = 4). In each experiment, responses were first obtained during a baseline period with 10 s inter-train intervals (yellow), then during a test period with 2 s intervals (green), and finally after the return of the inter-train interval to 10 s (orange). Each data point indicates the average value of 5 consecutive trains. Grey and black curves represent exponential fits to the control and Rose Bengal data, with time constants of 19 trains and 9 trains, respectively. **B**: Similar to A but for $s_1$. The time constants of exponential fits are 19 trains for control and 5 trains for Rose Bengal. **C**: Similar to A but for $s_2$. The time constants of exponential fits are 16 trains for control and 10 trains for Rose Bengal. **D**: Similar to A but for $S_{5-8}$. The time constants of exponential fits are 22 trains for control and 9 trains for Rose Bengal.

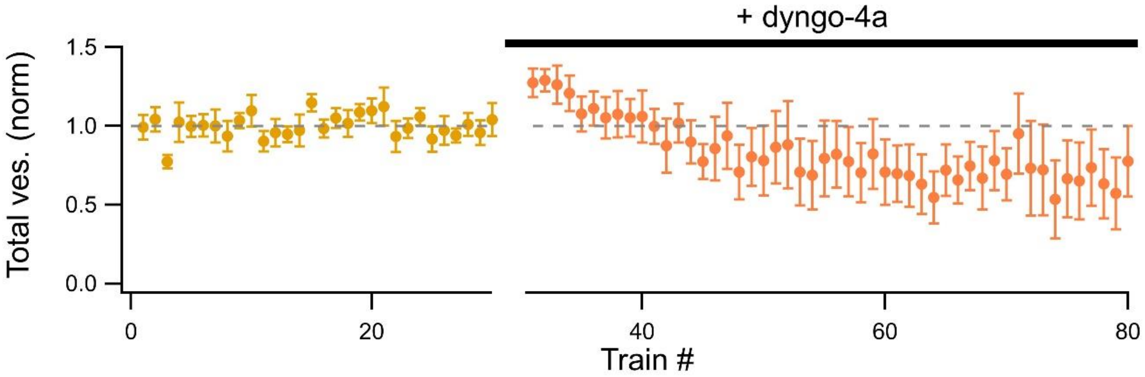


**Supplementary Figure 4: Dyngo-4a did not have a strong effect on SV release with 10 s inter-train intervals**

Group results showing the effect of dyngo-4a (30 µM; >20 min bath application) on the total SV release per train when the inter-train interval was kept at 10 s.

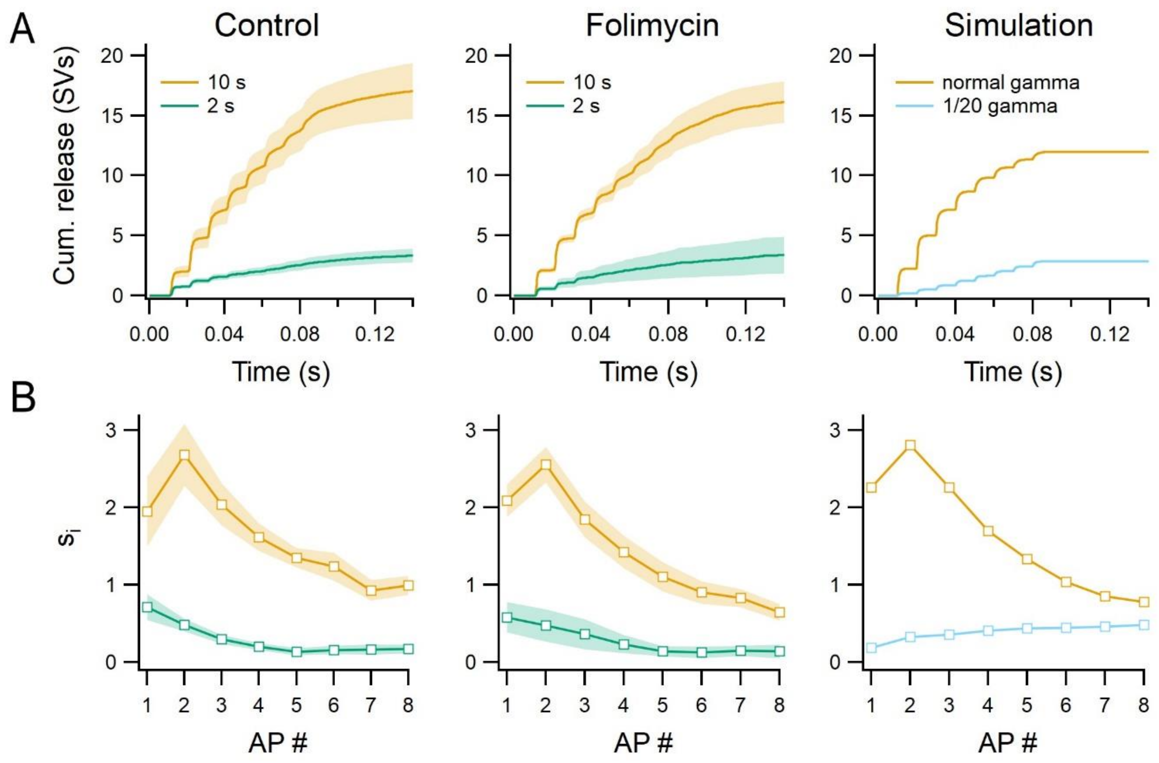


**Supplementary Figure 5: Comparison of SV release pattern between control and folimycin experiments, and simulation.**

**A:** Cumulative SV release during baseline stimulation (with 10 s intervals; yellow) and during the last 30 trains with 2 s intervals (green) in control experiments (first column; n = 9) and in experiments with folimycin (middle column; n = 7). Last column: Monte Carlo simulation was performed as described in the Supplementary Method. To mimic a reduction in release probability, the values of gamma were decreased by 20-fold (blue trace) compared to control (yellow trace). **B:** The corresponding number of SVs released per AP ($s_i$) as a function of AP number.

**Supplementary Methods**

**Simulation of SV release**

Monte Carlo simulation of SV release (**Suppl. Fig. 5**) were performed as previously described (Miki et al., 2018; Tran et al., 2022). Parameter values are shown in Table 1. Simulation of presynaptic AP-evoked $Ca^{2+}$ transients were done in CalC (version 7.9.4; Matveev et al., 2002), as described previously (Miki et al., 2018; Tran et al., 2022). The single-channel current had an amplitude of 0.2 pA and a Gaussian distribution with a half-width of 0.47. To simulate SV release for the decreased p scenario, the values of $\gamma$ for both normal and slow release rates were reduced by 20-fold. Calculations were done 3000 times for each condition and the averaged values are displayed.

**Table 1: Parameter values for simulation of SV release**

| **Simulation parameter** | **Value** | **Unit** | **Reference** |
|---|---|---|---|
| Number of docking sites (N) | 5 | | Determined experimentally by variance-mean analysis (Malagon et al., 2016) |
| Docking site occupancy ($\delta$) | 0.47 | | (Malagon et al., 2020) |
| Replacement site occupancy | 0.9 | | |
| Allosteric model for normal release rate | | | |
| $k_{on}$ | $5 \times 10^8$ | $M^{-1}s^{-1}$ | |
| $k_{off}$ | 5000 | $s^{-1}$ | |
| b | 0.75 | | |
| $\gamma$ | 2000 | $s^{-1}$ | |
| f | 31.3 | | |
| Allosteric model for slow release rate | | | |
| $k_{on}$ | $5 \times 10^7$ | $M^{-1}s^{-1}$ | |
| $k_{off}$ | 2000 | $s^{-1}$ | |
| b | 0.5 | | |
| $\gamma$ | 1200 | $s^{-1}$ | |
| f | 31.3 | | |
| Recovery time from slow to normal release rate | 150 | ms | |
| Recruitment from a replacement site to a docking site | | | |
| $V_{max}$ | 800 | $s^{-1}$ | |
| $K_d$ | 2 | µM | |
| Recruitment from the intermediate pool to a replacement site | | | |
| $V_{max}$ | 90 | $s^{-1}$ | |
| $K_d$ | 2 | µM | |
| Replenishment of each SV site in the intermediate pool | | | |

| | | | |
|---|---|---|---|
| $Ca^{2+}$ independent time constant | 0.35 | s | |
| $V_{max}$ | 30 | $s^{-1}$ | |
| $K_d$ | 5 | µM | |
| n | 5 | | |
| IP size at rest | 8 | SVs | (Tran et al., 2022) |
